# Metabolomics profiling reveals association of fermented herbal feed additive, growth performance and gut microbiota in piglets

**DOI:** 10.1101/2024.07.21.604508

**Authors:** Ruiming Xiao, Lingling Wang, Zhiqiang Tang, Xueqiao Qian, Jian Wang, Yingli Lian, Jiayi Tang, Jiarou Xu, Ying Lin, Baojun Shi, Pan Xu, Qiongsi Xiong

**Author notes:** Corresponding author: Baojun Shi, Animal Husbandry and Fisheries Research Center of Guangdong Haid Group Co., Ltd., Guangzhou 511400, China.

## Abstract

Fermented Chinese medicine (FCM), as a functional feed additive, has been widely recognized to play a significant role in protecting the intestinal health of piglets and enhancing productive performance. However, the relationship between the active components of FCM, gut microbiota, and their beneficial effects on animal performance remains unclear. In this study, metabolomics analysis revealed a significant increase in the main contents of lactic acid and propionic acid in FCM, while most glycosides and their derivatives decreased after three days of microbial fermentation. Subsequently, piglets were fed a basic diet supplemented with 1% FCM, while the control group received only the basic diet. The results indicated a significant increase in feed intake and average daily gain within 14 days (*P*<0.05) due to FCM supplementation. Additionally, FCM significantly improved feed efficiency from 1.76 to 1.50 (*P*<0.05). Meanwhile, piglets in the FCM group exhibited significantly lower frequencies of diarrhea and coughing, indicating better health. Moreover, high-throughput sequencing analysis revealed higher abundances of *Lactobacillus agilis*, *Megasphaera elsdenii*, *Ligilactobacillus*, and *Veillonellaceae* in piglets fed with FCM. In summary, FCM increased the content of active ingredients through microbial fermentation and regulated the intestinal microbiota to improve the health of piglets. FCM offers a promising potential for enhancing production performance and economic efficiency in the livestock industry.

## 1. Introduction

The management of weaning piglets is considered one of the most critical periods in pig breeding. Due to changes in environmental, social, and dietary stressors, a decrease in piglet feed intake and feed efficiency can result in symptoms such as diarrhea and infection, severely impacting livestock productivity [1, 2]. Approximately 15–20% of all pig born are estimated to die during the farrowing process or early lactation [3]. Recently, various substances have been used as feed additives to improve the growth performance of weaned piglets, including fermented probiotics [4], acidifiers [5], and plant-derived ingredients [6]. However, due to individual differences among animals, environmental variations, and unclear mechanisms, the effectiveness of these feed additives has shown significant variation.

Traditional Chinese medicine (TCM), a plant-based material exhibiting various physiological properties such as antibacterial, anti-inflammatory, and antioxidant effects, has been widely used for the healthcare of weaned piglets [7]. Recent studies have reported that traditional Chinese medicine or its active ingredients enhance immune function, digestion, and absorption, balance intestinal microecology, and effectively promote pig health and production performance [8, 9]. Furthermore, with the development of probiotics fermentation, it was confirmed that Lactobacillus fermentation of herb mixtures improved in-vitro anti-inflammatory activity and reduced blood endotoxin and CRP levels, as well as gut permeability [10]. Another study indicated that probiotic fermentation positively contributed to the release of functional compounds from Chinese herbs [11]. These active compounds can directly act on the host or exert their effects through secondary metabolic products generated by gut microbiota metabolism, such as short-chain fatty acids (SCFAs), amino acids, and vitamins [12, 13]. Nevertheless, TCM generally constitute a complex group, and the relationship between their core components and piglet growth performance has not been systematically elucidated, which may affect the efficacy of various formulations under different breeding environments.

In this research, fermented traditional Chinese medicine (FCM) was prepared using a probiotics mixture for fermentation. Bacterial counts, organic acids, and variations in active components were investigated using both targeted and untargeted metabolomics. Subsequently, FCM was incorporated into the basal diet of piglets as a feed additive to explore its impact on their growth performance. Average daily weight gain, feed intake, and feed efficiency were recorded, and changes in the intestinal microbiota of piglets during the feeding process were analyzed using high-throughput sequencing. T The study revealed the interactive patterns between changes in the core components of FCM and the growth performance and intestinal microbiota of piglets, providing a theoretical basis for applying FCM as a piglet feed additive.

## 2. Materials and Methods

### 2.1. Preparation of Fermented Chinese medicine feed additive

The complex probiotic powder that containing *Lactobacillus plantarum*, *Bacillus subtilis*, and *Saccharomyces cerevisiae* was already available from our laboratory. Following the formula in Table 1, the microbial powder and the materials were thoroughly mixed, ensuring that the initial water content of the fermentation was controlled at 40%. The mixture was then packed into breathable bags and compressed appropriately. Finally, the breathable bags were sealed and incubated at 37℃ for 3 days.

**Table 1.** Ingredient composition of Fermented Chinese medicine feed additive.

| Ingredients | FCM content (%) |
| --- | --- |
| Corn germ meal | 20 |
| Rapeseed meal | 25 |
| Corn Bran | 10 |
| Oily rice bran | 20 |
| Rice Husk Powder | 15 |
| Chinese herbal compounds <sup>1</sup> | 5 |
| Probiotic compounds | 5 |
| Total | 100 |

<sup>1</sup> The traditional Chinese medicine compound is a package containing herbs *Astragalus*, *Codonopsis pilosula*, *Glycyrrhiza*, and *Atractylodes*, prepared by our laboratory.

### 2.2. pH measurement and viable bacteria count

The 5 g of FCM was weighed and 95 ml of distilled water was added, then vortexed for 5 minutes to resuspend. The supernatant was then collected for pH measurement. Additionally, the bacterial supernatant was serially diluted and plated on MRS, LB, and PDA media respectively for counting lactic acid bacteria, Bacillus, and yeast.

### 2.3. Untargeted metabolomics analysis of the active components

The 100 mg of feed sample (before and after fermentation) was accurately weighed into a centrifuge tube. 1mL of extraction solvent (water/acetonitrile/isopropanol, 1:1:1, v/v/v) was added into the tube, followed by vortexing for 1 min and sonication at 4℃ for 30 min. Centrifuging was performed at 12000 rpm for 10 min at 4℃ to collect the supernatant. Then, the supernatant was collected and vacuum dried. The residue was resuspended in 200 μL of acetonitrile solution (50%, v/v). After vortexing, the sample was centrifuged at 14000 rpm at 4℃ for 15 min, and the resulting supernatant was used for instrumental analysis.

Data collection was carried out on Ultra-High Performance Liquid Chromatography (Vanquish, UPLC, Thermo, USA) equipped with a Waters HSS T3 column (100×2.1 mm, 1.8 μm) and High-Resolution Mass Spectrometer (Q Exactive HFX, Thermo, USA). The mobile phase consisted of Phase A (water solution of formic acid, 0.1%, v/v), and Phase B (acetonitrile solution of formic acid, 0.1%, v/v). Flow rate: 0.3 mL/min; Column temperature: 40℃; Injection volume: 2 μL. The gradient elution was programmed as follows: initially, Phase A/B was 100:0 (v/v) for 0-1 min, then shifted to 5:95 (v/v) at the 12 min and sustained for one minute, afterwards reverting to 100:0 (v/v) at 13 min and sustained until 17 min.

The ESI source operation parameters were as follows: source temperature 350 °C; ion spray voltage 3000 V (positive), −2800 V (negative); Sheath gas and curtain gas were set at 40 and 10 psi, respectively. The results were compared with the internal database of Sanshu Biotechnology Co., Ltd. (Nantong, China).

### 2.4. Quantification of organic acid content

The 0.1-0.5 g sample was added to a centrifuge tube and mixed with 1 mL of extraction solution (methanol/chloroform, 7:3, v/v). The mixture was then placed on ice for 30 minutes. Then, 0.6 mL of water was added and mixed, followed by centrifugation at 12,000 rpm for 10 minutes at 4℃. After collecting the supernatant, the extraction process was repeated twice. The supernatants from the two extractions were then combined and mixed for further use. Subsequently, 40 μL of the standard solution was added to the centrifuge tube, followed by 10 μL of 0.1 M EDC and 10 μL of 0.1 M 3NPH solution. The reaction mixture was then incubated at 40℃ for 30 minutes for derivatization. Detection was carried out on a UPLC-MS system equipped with a Waters BEH C18 column (50×2.1 mm, 1.8 μm). The instrument parameters were consistent with the method in section 2.2. The results were then compared with the internal database of Sanshu Biotechnology Co., Ltd. (Nantong, China).

### 2.5. Animals, Diets and Experimental Design

A total of 20 nursery piglets (Duroc × Landrace × Yorkshire) with similar body conditions and parities (body weight: 13.53 ± 0.74 kg, age: 44 ± 1 days) were randomly assigned to either the control group or the experimental group. The piglets in the control group were fed a basal diet, while those in the experimental group were fed a modified basal diet, in which the rice bran was replaced by FCM at an additive amount of 1%. The composition and nutrient levels of the two diets are listed in Table 2. All protocols in the study were approved by the Ethics Committee of the South China University of Technology Experimental Animal Center and followed the Regulations of Guangdong Province on the Administration of Experimental Animals (2019G041).

**Table 2.**
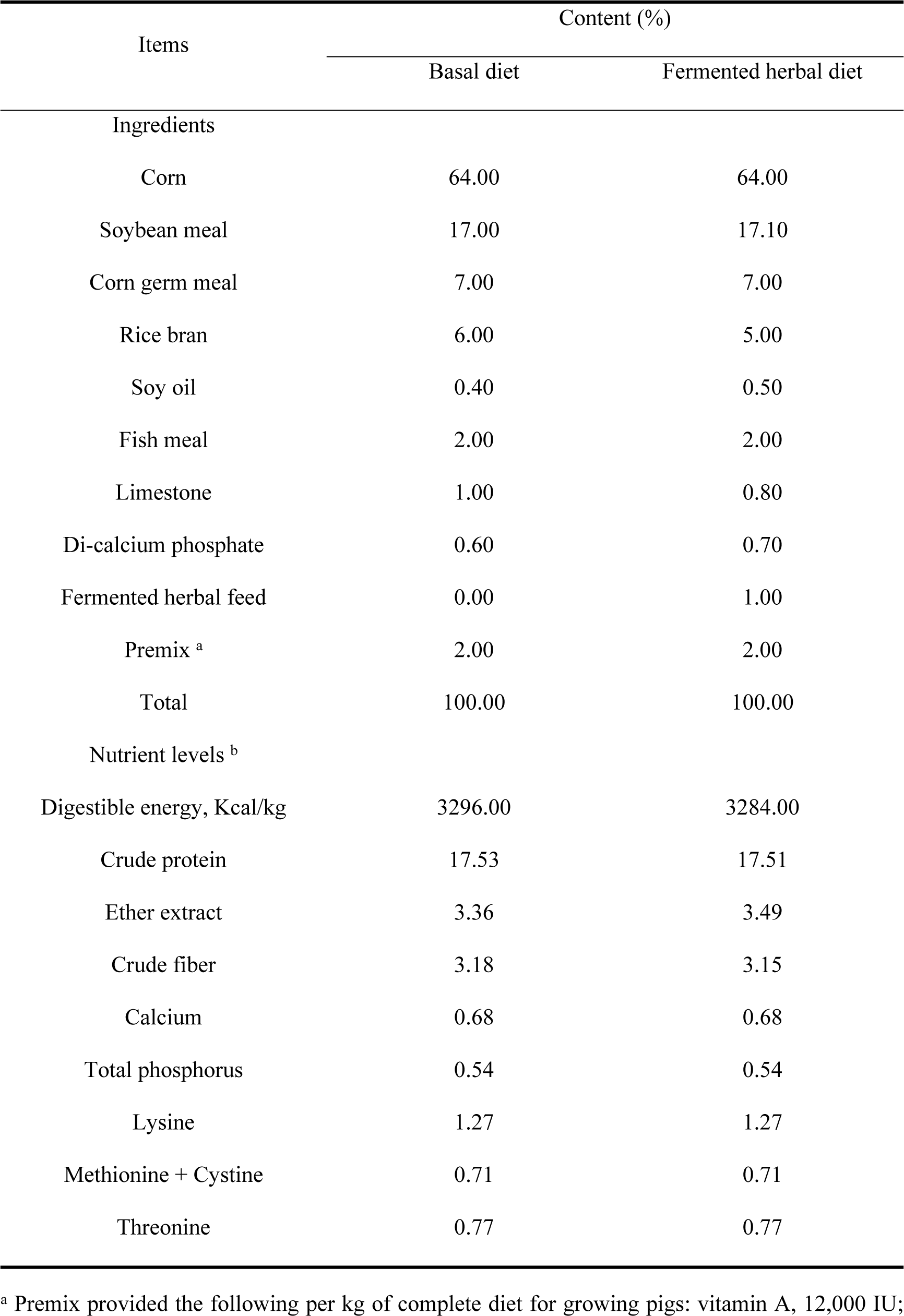
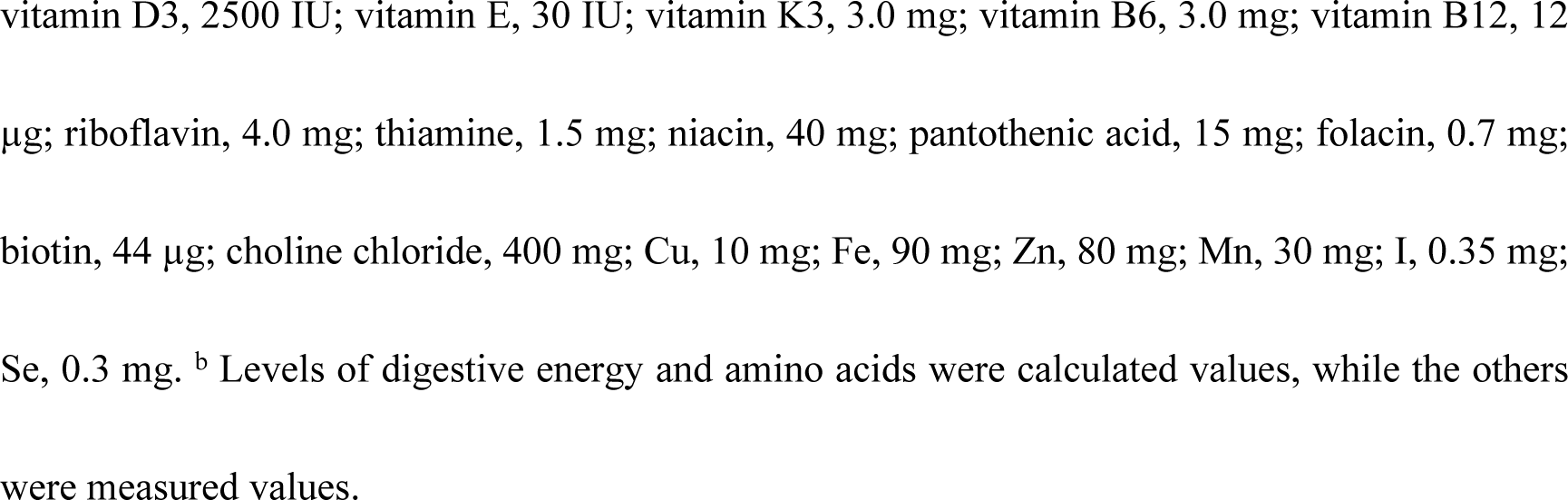
Composition and nutrient levels of the different diet (air-dry basis).

| Items | Content (%) |  |
| --- | --- | --- |
|  | Basal diet | Fermented herbal diet |
| Ingredients |  |  |
| Corn | 64.00 | 64.00 |
| Soybean meal | 17.00 | 17.10 |
| Corn germ meal | 7.00 | 7.00 |
| Rice bran | 6.00 | 5.00 |
| Soy oil | 0.40 | 0.50 |
| Fish meal | 2.00 | 2.00 |
| Limestone | 1.00 | 0.80 |
| Di-calcium phosphate | 0.60 | 0.70 |
| Fermented herbal feed | 0.00 | 1.00 |
| Premix <sup>a</sup> | 2.00 | 2.00 |
| Total | 100.00 | 100.00 |
| Nutrient levels <sup>b</sup> |  |  |
| Digestible energy, Kcal/kg | 3296.00 | 3284.00 |
| Crude protein | 17.53 | 17.51 |
| Ether extract | 3.36 | 3.49 |
| Crude fiber | 3.18 | 3.15 |
| Calcium | 0.68 | 0.68 |
| Total phosphorus | 0.54 | 0.54 |
| Lysine | 1.27 | 1.27 |
| Methionine + Cystine | 0.71 | 0.71 |
| Threonine | 0.77 | 0.77 |

^a^ Premix provided the following per kg of complete diet for growing pigs: vitamin A, 12,000 IU; vitamin D3, 2500 IU; vitamin E, 30 IU; vitamin K3, 3.0 mg; vitamin B6, 3.0 mg; vitamin B12, 12 µg; riboflavin, 4.0 mg; thiamine, 1.5 mg; niacin, 40 mg; pantothenic acid, 15 mg; folacin, 0.7 mg; biotin, 44 µg; choline chloride, 400 mg; Cu, 10 mg; Fe, 90 mg; Zn, 80 mg; Mn, 30 mg; I, 0.35 mg; Se, 0.3 mg. ^b^ Levels of digestive energy and amino acids were calculated values, while the others were measured values.

### 2.6. Feeding and Management

During the breeding process, FCM was mixed with the basal diet. Feeding was conducted three times per day, and piglets were provided ad libitum access to water and diet. The total daily feed intake of each group was recorded to calculate the average daily feed intake (ADFI), and the total weights were measured on days 0 and 14 to determine the average daily weight gain (ADG) and feed conversion ratio (FCR). Throughout the experiment, the incidence of diarrhea in piglets was continuously monitored, and the diarrhea rate (DR) was calculated using the following formula:

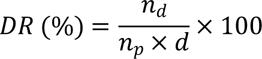

Annotation: *n_d_*, the number of piglet diarrhea during the experiment period; *n_p_*, the total number of piglets; *d*, the number of experimental days.

### 2.7. Fecal sample collection

Fecal samples from the piglets were collected on days 0, 4, and 14 to measure the variation in gut microbiota before and after feeding with different diets. The defecation of the piglets was monitored, and the fecal samples were collected promptly. The fecal samples were then transferred under dry ice conditions, and genome extraction and sequencing were conducted by Novogene Co., Ltd. (Tianjin, China).

### 2.8. Next generation sequencing and bioinformatics analysis

The total bacterial genomic DNA was used as template to amplify the V3-V4 hypervariable region of the 16S rRNA gene using primers 341 F (5′-CCTAYGGGRBGCASCAG-3′) and 806 R (5′-GGACTACHVGGGTWTCTAAT-3′). The sequencing libraries were quantified using Qubit and real-time PCR, and size distribution was assessed with a bioanalyzer. The quantified libraries were pooled and sequenced on the Illumina HiSeq 2500 platform to generate 2 × 250 bp paired-end reads. Quality control and clustering results were analyzed using QIIME 2.0 software. Optimized sequences were clustered at a 97% similarity threshold and blasted against the Silva Database (https://www.arb-silva.de) to identify the taxa. Further analyses, including alpha diversity, beta diversity, and taxonomic distinctness, were conducted in R (version 4.2.1).

### 2.9. Statistical Analysis

All results were analyzed statistically using GraphPad Prism 7.0 software. The Student’s t-test was used to assess the significance between paired results, while the Kruskal-Wallis test was used to compare ranks for non-paired microbial abundance results. A P-value of less than 0.05 was considered statistically significant.

## 3. Results

### 3.1. Variation of the active components in FCM feed

After 3 days of microbial fermentation, the active compounds in FCM feed were revealed using a non-targeted metabolomics method. According to Fig. 1A, the composition of active constituents was significantly modified by fermentation. A total of 1,029 herbal metabolites were identified by UPLC-MS, with 53 and 19 compounds individually labeled in pre- and post-fermentation, respectively. Meanwhile, the variation of 957 common metabolites was shown in a volcano plot as presented in Fig. 2B. It was found that 231 compounds were distinctly consumed and 192 were significantly increased after fermentation. Then, the extremely significant (*P*<0.001) variations were counted, as presented in Fig. 1C. The microbial fermentation enriched 29 active compounds, among which xylotetraose, 3-methyluridine, crotananine, virosine B, isonormangostin, and 10-deacetylbaccatin were the most abundant features. Another 79 compounds decreased in FCM feed, most of which were derivatives of glycosides and phenols, such as emodin-8-glucoside, guanosine, skimming, caffeic acid-4-O-glucuronide, and 10-O-trans-p-coumaroylscandoside, etc.

**Figure 1.**
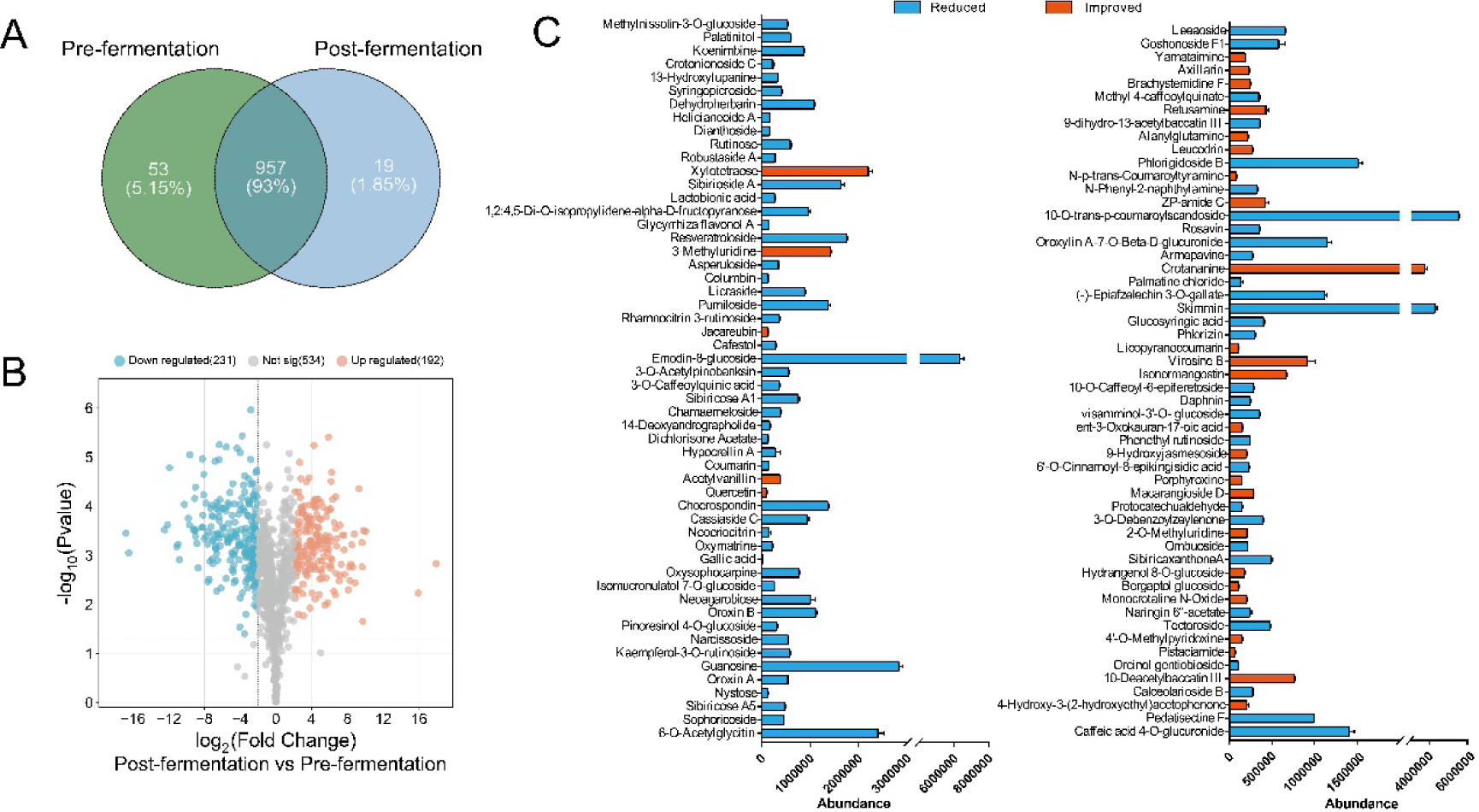
Variation of active components in FCM after fermentation of probiotics. (A) Venn diagram of total variation features; (B) Volcano plot of significantly changed active components; (C) Barplot of totally consumed and appeared active components.

**Figure 2.**
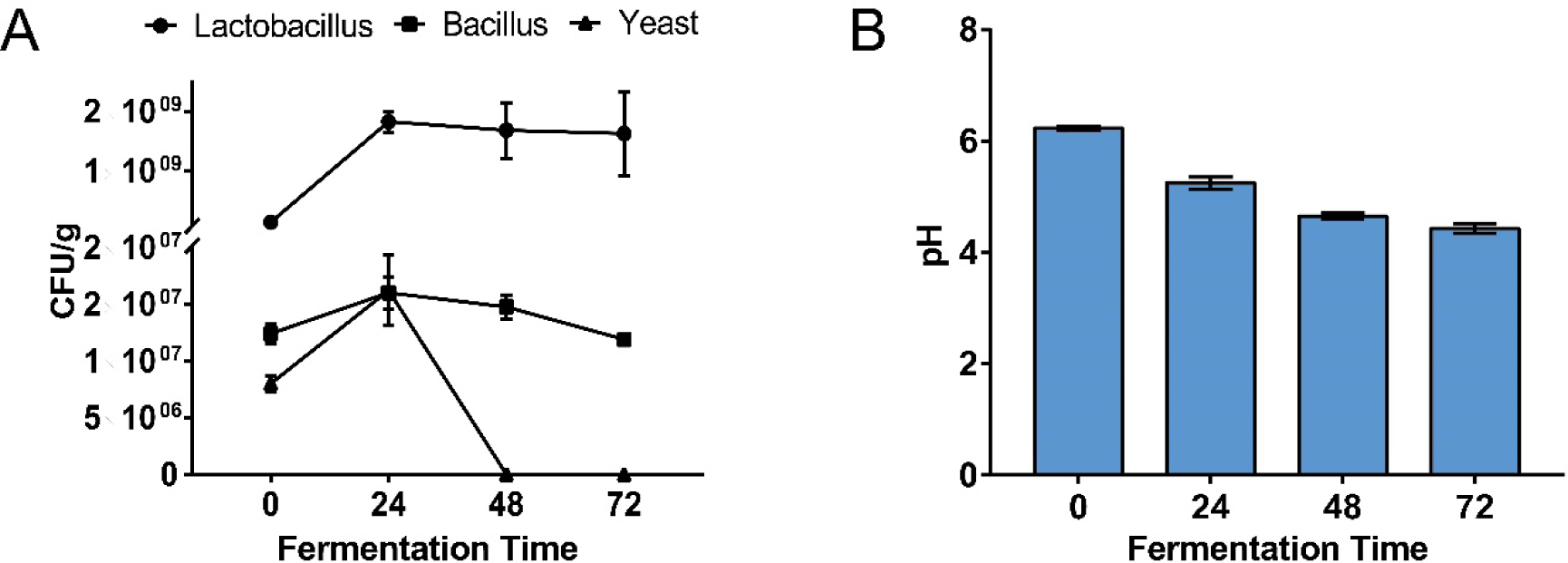
Variations of (A) viable bacteria counts and (B) pH of FCM during 72-h fermentation.

### 3.2. Variation of microbial and organic acids content in FCM feed

During the fermentation period, the active constituents in FCM were released, and the beneficial bacteria proliferated by utilizing the nutrients in FCM. According to Fig.2A, after 24 h of fermentation, Lactobacillus was the most abundant group, with its number increasing to 2.0 ×1010 CFU/g and remaining relatively stable at 48 and 72 h. Besides, the number of Bacillus showed little change during the 72-h fermentation (∼108 CFU/g), while yeast increased in the first 24 h and decreased in the next 48 h (∼104 CFU/g). As the lactic acid bacteria grew, the microbial fermentation also resulted in a decrease in pH, with the final pH reaching 4.43 at 72 h (Fig.2B). To further reveal the composition of organic acids, a targeted metabolomics method was applied to measure their concentrations (Table 3.). Obviously, lactic acid was the most abundant, showing a 100-fold increase compared to pre-fermentation levels. Additionally, propionic acid increased by 268-fold (from 88.15 to 23,733.84 μg/L), while citric acid and malic acid decreased to 29,240.99 and 342.28 μg/L, respectively. There were also varying degrees of changes among other metabolites such as isobutyric acid (*P*=0.0411), butyric acid (*P*=0.0055), valeric acid (*P*=0.0150), fumaric acid (*P*<0.0001), maleic acid (*P*=0.0014), and succinic acid (*P*=0.1354). However, their concentrations were significantly lower than those of the abundant acids; therefore, the less abundant acids were excluded from the discussion.

**Table 3.** Variation of organic acids in feed after supplementation of FCM.

| Organic acid | Concentration (µg/L) |  | *P*-value |
| --- | --- | --- | --- |
|  | Pre-fermentation | Post-fermentation |  |
| Lactic acid | 2121.01 ± 71.27 | 237991.03 ± 18638.48 | 0.0031 |
| Propionic acid | 88.15 ± 7.66 | 23733.84 ± 2364.88 | 0.0050 |
| Isobutyric acid | 35.64 ± 0.64 | 218.13 ± 54.02 | 0.0411 |
| Butyric Acid | 23.15 ± 1.04 | 209.22 ± 19.51 | 0.0055 |
| Malic acid | 20856.31 ± 443.37 | 342.28 ± 2.06 | 0.0002 |
| Succinic acid | 3246.64 ± 97.26 | 6753.35 ± 2035.34 | 0.1354 |
| Valeric acid | 6.04 ± 0.18 | 185.75 ± 31.47 | 0.0150 |
| Fumaric acid | 2678.25 ± 20.68 | 3.22 ± 0.10 | < 0.0001 |
| Citric acid | 41089.32 ± 304.31 | 29240.99 ± 765.74 | 0.0024 |
| Maleic acid | 2265.22 ± 23.43 | 163.60 ± 109.25 | 0.0014 |
| Malonic acid | 2370.71 ± 115.37 | 2218.23 ± 1218.43 | 0.8764 |

### 3.3. Effects of FCM feed on growth performance of weaned piglets

The effects of the different feeding diet on the performance of weaned pigs are shown in Table 4. Weaned pigs in the FCM+basal diet group had a higher average daily gain (ADG) from days 0 to 14 compared to those in the basal diet (*P*<0.01). Although no significant differences in the average daily feed intake (ADFI) of weaned pigs from days 0 to 14 were observed among different dietary treatments (*P*=0.1744), a significant decrease in the feed conversion ratio (FCR) was found in the Basal diet+FCM group, from 1.76 to 1.50 (*P*<0.0001). Additionally, the addition of FCM to the basal diet improved disease resistance. The diarrhea rate decreased from 3.57% to 0.36%, and the cough frequency reduced from 13 to 7 during the feeding period. These results indicate that the addition of FCM improved growth performance, feed efficiency, and overall health condition.

**Table 4.** Effects of additive FCM in basal diet on growth performance of weaned piglet.

|  | Pre-<br>weight | Post-<br>weight | ADG | ADFI | FCR | Diarrhea<br>rate | Cough<br>frequency |
| --- | --- | --- | --- | --- | --- | --- | --- |
| Basal diet | 14.23<br>± 0.25 | 22.98 ±<br>0.11 | 0.63 ±<br>0.03 | 1.10 ±<br>0.12 | 1.76 ±<br>0.12 | 3.57 <sup>a</sup> | 13 <sup>b</sup> |
| Basal<br>diet+FCM | 14.15<br>± 0.16 | 23.90 ±<br>2.62 | 0.70 ±<br>0.07 | 1.04 ±<br>0.06 | 1.50 ±<br>0.06 | 0.36 <sup>a</sup> | 7 <sup>b</sup> |
| *P* value | 0.4052 | 0.2818 | 0.0094 | 0.1744 | < 0.0001 | - | - |

<sup>a,b</sup> The diarrhea rate and cough frequency was be counted according the entire breeding period without subfield.

### 3.4. Effects of FCM feed on gut microbial composition of Weaned piglets

The extracted DNA was employed to 16S rRNA sequencing, targeting the V3-V4 region, to analyze the composition of the gut microbiota in different feed groups. The α-diversity indices served as crucial indicators reflecting the variation in the overall microbiota composition of the samples (Table 5). In comparison to the baseline, the Chao1, observed features, Shannon, and Simpson indices of the basal diet and basal diet+FCM groups showed no significant variation (*P*>0.05). Furthermore, the Venn diagram (Fig. 3A) and principal component analysis (PCA) (Fig. 3B) revealed that the composition of the gut microbiota was significantly modified by different feeding diets. Hence, the linear discriminant analysis effect size (LEfSe) was conducted to distinguish the dominant taxa in different feeding groups (Fig. 3C). The main phyla of the gut microbiota were *Bacteroidetes*, *Firmicutes*, and *Proteobacteria*, which accounted for 98% of the bacterial abundance in the gut of piglets. Compared with the baseline, the Basal diet+FCM significantly increased the abundances of *Megasphaera*, *Ligilactobacillus*, *Lactobacillus*, and *Veillonellaceae* (*P*<0.05), while the basal diet only enriched *Prevotella* and *Alloprevotella* (*P*<0.05). Considering the FCM contained a high amount of *Lactobacillus*, the enrichment of this taxon in the basal diet+FCM group might be related to the intake of these bacteria.

**Figure 3.**
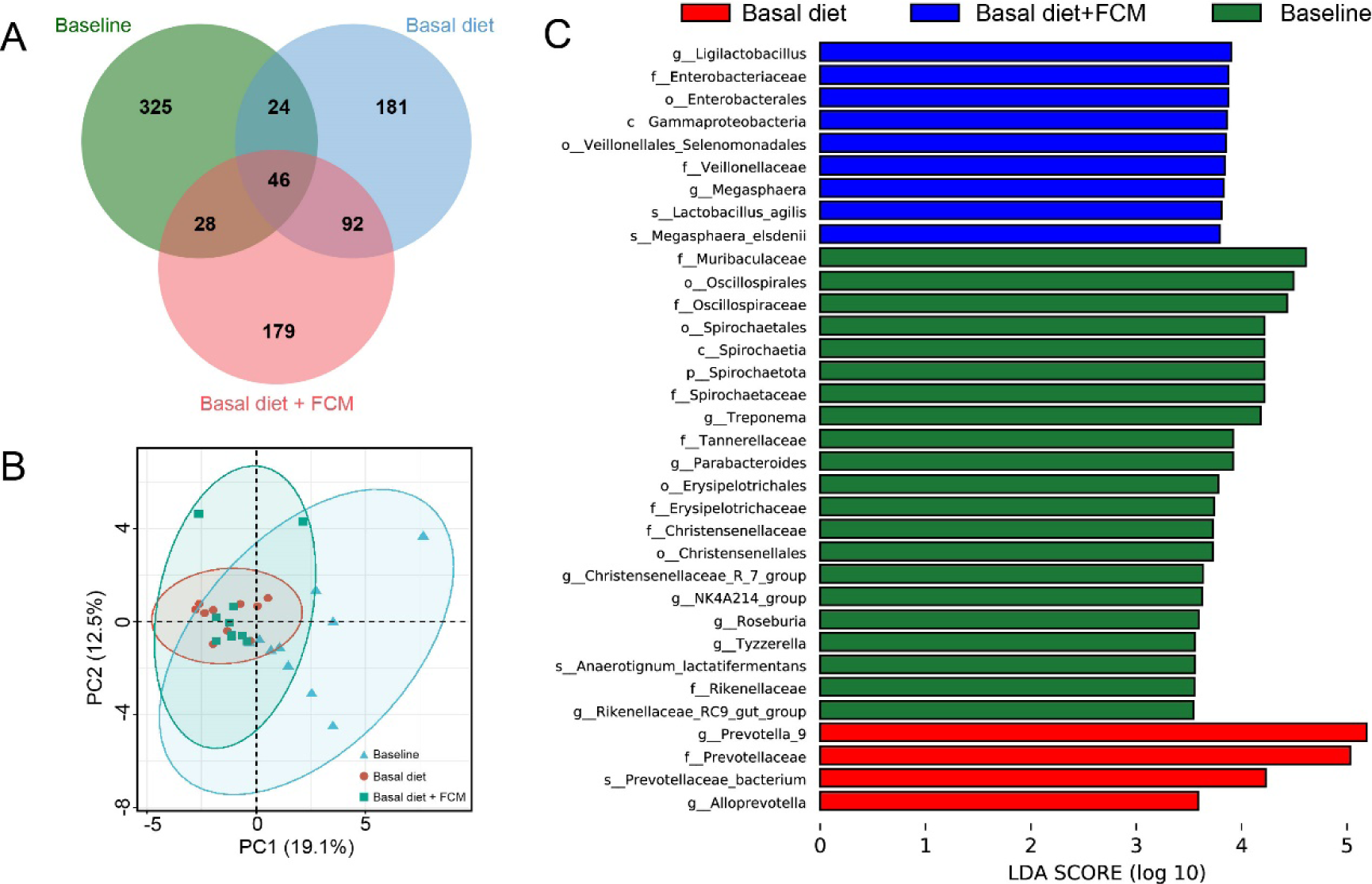
Variations of gut microbiota in (A) Venn diagram of features; (B) principal component analysis; (C) LEfSe analysis of gut microbiota in different feeding group.

**Table 5.** Effects of additive FCM in basal diet on alpha-diversity index of gut microbiota.

|  | Chao1 | Observed features | Shannon | Simpson |
| --- | --- | --- | --- | --- |
| Baseline | 478.81 ± 90.46 | 460.00 ± 85.20 | 6.39 ± 0.35 | 96.94 ± 1.06 × 10<sup>-2</sup> |
| Basal diet | 478.02 ± 100.98 | 461.10 ± 97.80 | 6.52 ± 0.48 | 97.47 ± 0.80 × 10<sup>-2</sup> |
| Basal diet + FCM | 456.56 ± 90.46 | 441.40 ± 70.14 | 6.46 ± 0.27 | 97.12 ± 0.82 × 10<sup>-2</sup> |

## 4. Discussion

The active ingredients in traditional Chinese medicine have been verified to have various functions, such as improving immune response, maintaining gut barrier integrity, and exhibiting antioxidant properties. Microbial fermentation is also an effective approach to enhance the performance of traditional Chinese medicine. It has been reported that the total free flavonoid content in *Pericarpium Citri Reticulatae* increased by 48.12% after fermentation by probiotics [14]. Additionally, Chen et al. supplemented herbal medicine into the daily diet of piglets, resulting in significant improvment in average daily gain (ADG) and average daily feed intake (ADFI), as well as gut microbial diversity [15].

According to statistics, over 70% of the fully consumed active ingredients are glycosides and their derivatives (Fig 1C). For example, three of the most abundant compounds, emodin-glucoside, 10-O-trans-p-coumaroylscandoside, and skimmin, were completely consumed after fermentation. However, due to the complexity of microbial metabolic pathways and the various active contents, we did not track the metabolites of these compounds. Hydrolysis reactions, commonly found in various microorganisms [16], were identified in the metabolism of active constituents. For instance, quercetin, an aglycone of flavonoid glycosides, significantly increases after microbial fermentation, indicating that probiotics hydrolyze flavonoid glycosides and release the aglycone. Our previous study has shown that high-level glycosides were hydrolyzed into low-level glycosides or aglycones, which might exhibit stronger biological activity and increased bioavailability [17]. Consequently, it was speculated that the health benefits of FCM might be enhanced by probiotics through the release of more functional active components. Furthermore, certain prebiotic oligosaccharides, such as xylotetraose, were significantly enriched after fermentation. According to previous reports, a xylanase derived from *Bacillus paralicheniformis* helped hydrolyze xylan and increased the content of xylobiose, xylotriose, and xylotetraose [18]. Meanwhile, these oligosaccharides can be utilized by probiotics or gut microbiota for proliferation and the production of beneficial secondary metabolites, such as organic acids, amino acids, and vitamins [19]. However, excessive microbial fermentation may also lead to the loss of nutritional and functional components. Therefore, the inoculation amount and fermentation time of FCM should be controlled to determine the optimal conditions for achieving the best concentration of active compounds.

During the fermentation process, we continuously monitored the acidity of FCM inoculated with three bacterial strains. Within the first 0-24 hours, the anaerobic fermentation occurring inside the FCM led to the rapid proliferation of lactic acid bacteria, while the aerobic Bacillus multiplied more slowly. Although the numbers of lactic acid bacteria and Bacillus tended to stabilize between 48-72 hours, the yeast population sharply declined at the 48-hour mark, possibly due to the production of antifungal metabolites in the FCM. Additionally, due to the continuous acid production by lactic acid bacteria, the pH consistently decreased and approached stability after 48 hours. Moreover, targeted metabolomics of organic acids revealed that lactic acid and propionic acid were the two main contributors to the pH decrease.

It has been confirmed that *Lacticaseibacillus rhamnosus* and *Limosilactobacillus reuteri* possess pathways for the synthesis of lactic and propionic acids [20]. However, the synthesis pathways for other short-chain fatty acids such as butyric acid, valeric acid, and isobutyric acid are rarely reported in this genus. Some studies suggest that lactic acid bacteria convert malic acid to lactic acid through malolactic fermentation [21], which is consistent with our findings that the abundance of malic acid significantly decreased, accompanied by a partial reduction in citric acid content. As feed acidifiers, lactic acid and propionic acid are also used to reduce the acid-binding capacity of feed and promote growth performance [22]. Furthermore, active probiotics can produce various cellular components such as enzymes, teichoic acids, and polysaccharides [23, 24], which regulate the immunity and gut microecology of piglets. Therefore, adding active probiotics may be more effective than directly supplementing with acidifiers.

Further breeding experiments indicated that the additive of FCM positively impacts the growth performance of piglets. Active components such as flavonoids, polysaccharides, and phenolic acids have been proven to enhance piglet feed intake. For instance, supplementation with ferulic acid was found to significantly increase liver SOD activity and improve lipid metabolism [25]. The additive of vanillic acid significantly improved ADG and ADFI, and enhanced gut barrier morphology in lipopolysaccharide-challenged piglets [26]. Hence, the effect of FCM might originate from the metabolism of complex carbohydrates by microorganisms, which produce secondary metabolites with immune-regulating properties. Moreover, probiotic fermentation not only increased the content of active components in FCM but also led to the substantial proliferation of lactic acid bacteria, thereby improving the acidity of the feed and providing sufficient gut probiotics for enhanced feed digestibility.

Due to the fibrous components in the basal diet, the abundance of *Bacteroidetes*, particularly *Prevotella*, was enriched in both the Basal diet and Basal diet + FCM groups, with the latter exhibiting less significant variation. Previous studies have indicated that *Prevotella* is closely associated with carbohydrate metabolism in piglets [27]. The composition and abundance of gut microbiota are constantly changing, especially during the early life of piglets [28]. Therefore, the baseline gut microbiota was also measured as a starting point reference. It was observed that xylan, one of the major fibrous components in herbal medicine [29], was closely related to the enrichment of *Prevotella* [30]. In addition, *Lactobacillus agilis* and *Ligilactobacillus* were also enriched by the FCM additive, which is closely related to the intake of active lactic acid bacteria in FCM. However, since this study focused on the effects of FCM on growth performance and gut microbiota regulation, we did not track the colonization of lactic acid bacteria after the 14-day breeding period.

Moreover, *Megasphaera elsdenii* and *Veillonellaceae* were significantly enriched by additive FCM. *Megasphaera elsdenii*, an anaerobic lactic acid-fermenting bacterium, has been reported to effectively restore the atrophy of the small and large intestinal mucosa when orally administered to weaned piglets [31]. The enrichment of *Veillonellaceae* was also observed in weaned piglets fed natural plant extracts, such as essential oils [32]. According to previous reports, the enrichment of beneficial taxa can directly result from the inhibition of potential pathogens such as *Escherichia coli* or the provision of substrates for commensal bacteria via “cross-feeding” [33]. However, the interaction between gut microbiota and active components in FCM is complex, and the relationships between specific taxa and active components remain to be elucidated. Additionally, due to restrictions in testing conditions and production environments, we did not evaluate the differences in growth performance between unfermented traditional Chinese medicine and FCM in piglets. This is a limitation of our study. Our future work will explore the functional components of FCM and their role in enhancing piglet immune function, particularly through the application of probiotics and postbiotics.

## 5. Conclusions

In conclusion, microbial fermentation significantly altered the composition of active components, increased the probiotic count, and enhanced the feed’s acidity by consuming the glycoside in FCM. As a functional feed additive, FCM significantly improved the ADG of piglets, reduced the FCR, and increased feed efficiency. It also lowered the diarrhea rate and cough frequency, thereby improving the health status of the piglets. Intestinal microbiota analysis showed that FCM enriched beneficial gut bacteria such as *Lactobacillus*, *Megasphaera*, and *Veillonellaceae*, demonstrating excellent effects on improving the growth performance and gut microbiota of piglets. In summary, FCM provides a viable solution to improve the growth performance and health status of piglets, showing strong potential for commercial application.

## Author Contributions

Conceptualization, R.X. and L.W.; methodology, L.W.; software, R.X.; validation, Z.T., J.X. and P.X.; formal analysis, R.X.; investigation, Y.L.; resources, X.Q. and J.W.; data curation, Q.X.; writing—original draft preparation, R.X.; writing—review and editing, J.T.; visualization, L.W. and Z.T.; supervision, Y.L.; project administration, B.S.; All authors have read and agreed to the published version of the manuscript.

## Funding

This research received no external funding.

## Data Availability Statement

The data that support the findings of this study are available from the corresponding author upon reasonable request.

## Conflicts of Interest

The authors declare no conflicts of interest.

## Notes

### Competing Interest Statement

The authors have declared that no competing interests exist.

